# “Landscape of 8q24.3-Encoded microRNAs and Their Prognostic Impact in Ovarian Cancer”

**DOI:** 10.64898/2026.03.06.710032

**Authors:** Kamil Filipek, Ivan Merelli, Federica Chiappori, Marianna Penzo

## Abstract

Ovarian cancer is the most lethal gynecological malignancy, largely because of late diagnosis and marked genomic instability, with high-grade serous ovarian cancer (HGSOC) representing its most common and aggressive subtype. Amplification of chromosome 8q24.3 is a recurrent event in HGSOC, yet the regulation and clinical relevance of the non-coding RNA output from this locus remain poorly defined. Here, we performed an integrative analysis of 8q24.3-encoded miRNAs in ovarian cancer using copy-number, transcriptomic, isoform-resolved, and clinical data from TCGA and NCBI datasets. We identified pronounced heterogeneity in miRNA abundance and strand usage across this locus. Copy-number gain broadly associated with increased miRNA expression, although this effect was not uniform across all candidates. Intronic miRNAs showed variable coupling with their host genes, indicating that mature miRNA output is shaped by both genomic dosage and post-transcriptional regulation. Isoform-level analysis revealed marked strand asymmetry and regulatory complexity, but did not strengthen copy-number or histotype associations compared with total miRNA measurements. Clinically, higher expression of miR-937, miR-4664, and miR-6849 was associated with improved overall survival in HGSOC. Functional enrichment of validated targets highlighted pathways related to cellular stress responses, senescence, p53 signaling, endocytosis, and metabolic adaptation. Together, these findings define 8q24.3 as a heterogeneous non-coding regulatory hub in ovarian cancer and provide a basis for future mechanistic and biomarker studies.

## 1. Introduction

Ovarian cancer (OC) is the most lethal gynecological malignancy worldwide, with more than 324,000 new cases and 207,000 deaths reported in 2022, ranking it eighth in both incidence and mortality across all female tumor types [1]. The high mortality to incidence ratio reflects the fact that most patients present with advanced disease, where the 5-year survival rate is ∼30%, compared to >90% when diagnosed early [2]. HGSOC is the predominant histological subtype of epithelial ovarian cancer (EOC), accounting for most cases, and is characterized by genomic instability, TP53 mutations, and copy number alterations [3–5]. Although advances in cytoreductive surgery, platinum-based chemotherapy, and targeted agents such as PARP inhibitors have improved progression-free survival, long-term survival remains poor [6,7], highlighting the need for new molecular insights and therapeutic opportunities.

miRNAs are small non-coding RNAs, typically 19-22 nucleotides in length, that regulate gene expression post-transcriptionally by guiding RNA-induced silencing complexes to the 3’UTR or 5’UTR of target mRNAs, leading to reduced or enhanced protein expression, respectively [8–10]. By simultaneously modulating multiple targets, miRNAs act as network-level regulators of cellular processes, including proliferation, apoptosis, and metastasis [11]. In ovarian cancer, widespread miRNA dysregulation has been documented and linked to tumorigenesis, metastasis, and therapy response, underscoring their relevance as biomarkers and potential therapeutic targets [12]. However, the functional impact of miRNAs is strongly influenced by their genomic context, such as transcriptional regulation and post-transcriptional processing [13].

Genomic loci recurrently altered in cancer often encode clusters of miRNAs whose expression and function are shaped by copy number changes [14], host gene regulation [15], epigenetic modification [16], and isoform diversity [17]. The chromosomal region 8q24.3 represents one such locus, harboring a dense repertoire of miRNA genes embedded within a complex genomic architecture. Individual miRNAs encoded at this locus have been implicated in cancer-associated pathways such as epithelial to mesenchymal transition (EMT) [18], stress-adaptive responses [19,20], PI3K/Akt/mTOR signaling [21,22], and autophagy [23,24]. Despite these observations, existing studies have largely focused on single miRNAs in isolation, providing limited insight into how the 8q24.3 locus operates as an integrated regulatory unit. In ovarian cancer, in particular, a systematic understanding of the expression landscape, regulatory dependencies, and transcriptional organization of 8q24.3-encoded miRNAs is currently lacking.

In this study, we present a comprehensive, multi-layered characterization of miRNAs encoded at the 8q24.3 locus in ovarian cancer by integrating genomic, transcriptional, and isoform-resolved analyses across TCGA and NCBI datasets. We define the genomic organization of all miRNAs within this region, quantify their expression patterns across HGSOC and non-serous EOC subtypes, assess the contribution of copy number alterations to their deregulation, and evaluate transcriptional coupling between intragenic miRNAs and their protein-coding host genes. By combining locus-level genomics with miRNA isoform profiling, our work provides the first integrative view of how amplification, host-gene context, and miRNA processing collectively shape the non-coding RNA output of the 8q24.3 region, offering new mechanistic insights and potential clinical relevance for ovarian cancer biology.

## 2. Material and Methods

### 2.1 Accession of the public database

The TCGA PanCancer Atlas database [3] deposited in cBioPortal [25] (accession on 21/02/2026) was used to acquire the chromosomal alterations, miRNAs genetic amplification percentage, miRNAs and host gene copy number, host mRNA expression values, MYC amplification and protein expression in ovarian serous cystadenocarcinoma tissues. Patient’s age at diagnosis, ethnicity, FIGO stage, mean tumor dimension, tumor residual disease, and type of chemotherapy treatment were downloaded from <u>portal.gdc.cancer.gov/projects/TCGA-OV</u> (accession on 21/02/2026). The TCGA database deposited in DIANA-microRNA Tissue Expression Database (miTED) [26] was used to acquire the miRNA expression values in ovarian serous cystadenocarcinoma tissues.

The GEO: GSE169314 database (accession on 21/02/2026), deposited in the National Center for Biotechnology Information (NCBI) Gene Expression, was used to analyze miRNA expression in different EOC subtypes.

### 2.2 Survival analysis

The association between the expression of 8q24.3-encoded miRNAs and overall survival (OS) in OC patients was performed using the publicly available KM Plotter tool (http://kmplot.com). Patient data from TCGA were analyzed, and expression levels of individual miRNAs were used to stratify patients into two groups based on the median expression value (high versus low expression). Survival curves were generated using the Kaplan–Meier method, and differences between groups were assessed with the log-rank test. Hazard ratios (HRs) with 95% confidence intervals (CIs) were calculated using Cox proportional hazards regression. Statistical significance was defined as *p* < 0.05.

### 2.3 miRNA target identification and enrichment

miRNA targets were identified using miRTarBase2025 v10.0 [27] (accession on 02/2026), selecting only experimentally validated targets, mainly by CLIP-Seq, Western blot, reporter assay, and qPCR. The resulting targets, 3230 genes, were used for Over-Representation Analysis (ORA) to identify significantly enriched pathways, biological processes, and gene collections among all targets. ORA analysis was conducted using ClusterProfiler (version 4.16.0) in R (version 4.5.1), and gene targets were annotated using GO (biological process and molecular function), KEGG, MSigDB (Hallmark and C6 genes), and Reactome databases. Only pathways with adjusted p-value <0.05 were considered.

### 2.4 Statistical analysis and data visualization

Expression and functional data were analyzed using a non-parametric statistical test Mann-Whitney U test. For >2 groups analyzed the p-values were calculated using Kruskal-Wallis test with post hoc Dunn test which were adjusted for multiple comparisons using Benjamini-Hochberg false rate correction (FDR). Correlation analyses were performed using Spearman’s rank correlation coefficient. All statistical analyses and data visualization were conducted in Python version 3.14.2 using standard scientific libraries, including scipy.stats [28] for statistical testing and correlation analysis, and statsmodels [29] for multiple testing correction. Data processing and visualization were performed using NumPy, Pandas, Matplotlib [30], and Seaborn. A *p* < 0.05 was considered statistically significant.

## 3. Results

### 3.1 Genomic landscape and annotation of 8q24.3-encoded miRNAs in OC

HGSOC shows a significant genomic instability, with recurrent copy number alterations that represent its primary characteristic [5,31]. Our analysis of a dataset of HGSOC from TCGA revealed that chromosomes 22 and 8 exhibited the highest frequency of alteration, exceeding 35% (Suppl. Fig. S1A). While chromosome 22 alterations were dominated by deletions (39%), chromosome 8 displayed a predominance of amplifications (23%) (Suppl. Fig. S1B). Given this amplification bias, we next focused on the 8q24.3 cytoband, a well-established cancer-associated locus that undergoes recurrent amplification [32] and harbors a dense cluster of microRNA genes. Notably, HGSOC exhibits the highest frequency of 8q24.3 amplification, whereas other tumor types, including breast, liver, and lung cancers show considerably lower amplification rates (Suppl. Fig. S1C). To thoroughly evaluate the role of 8q24.3-encoded miRNAs in OC, we initially mapped all 17 identified miRNAs within this region (Fig. 1A). Out of these, 14 miRNAs are intronic, two are intergenic, and one is exonic (Fig. 1B). It is important to note that copy number alterations for miR-1302-7 and miR-4472-1 were described in TCGA ovarian cancer samples (Suppl. Fig. S2A), however, these loci belong to multi-copy miRNA families (Suppl. Fig. S2B). In TCGA miRNAseq data, the expression of individual copies cannot be distinguished. Reads are assigned to the aggregate miR-1302 and miR-4472. Due to a lack of locus-specific expression values for miR-1302-7 and miR-4472-1, these miRNAs were not included in future studies. This left 15 candidates for expression profiling.

**Fig. 1.**
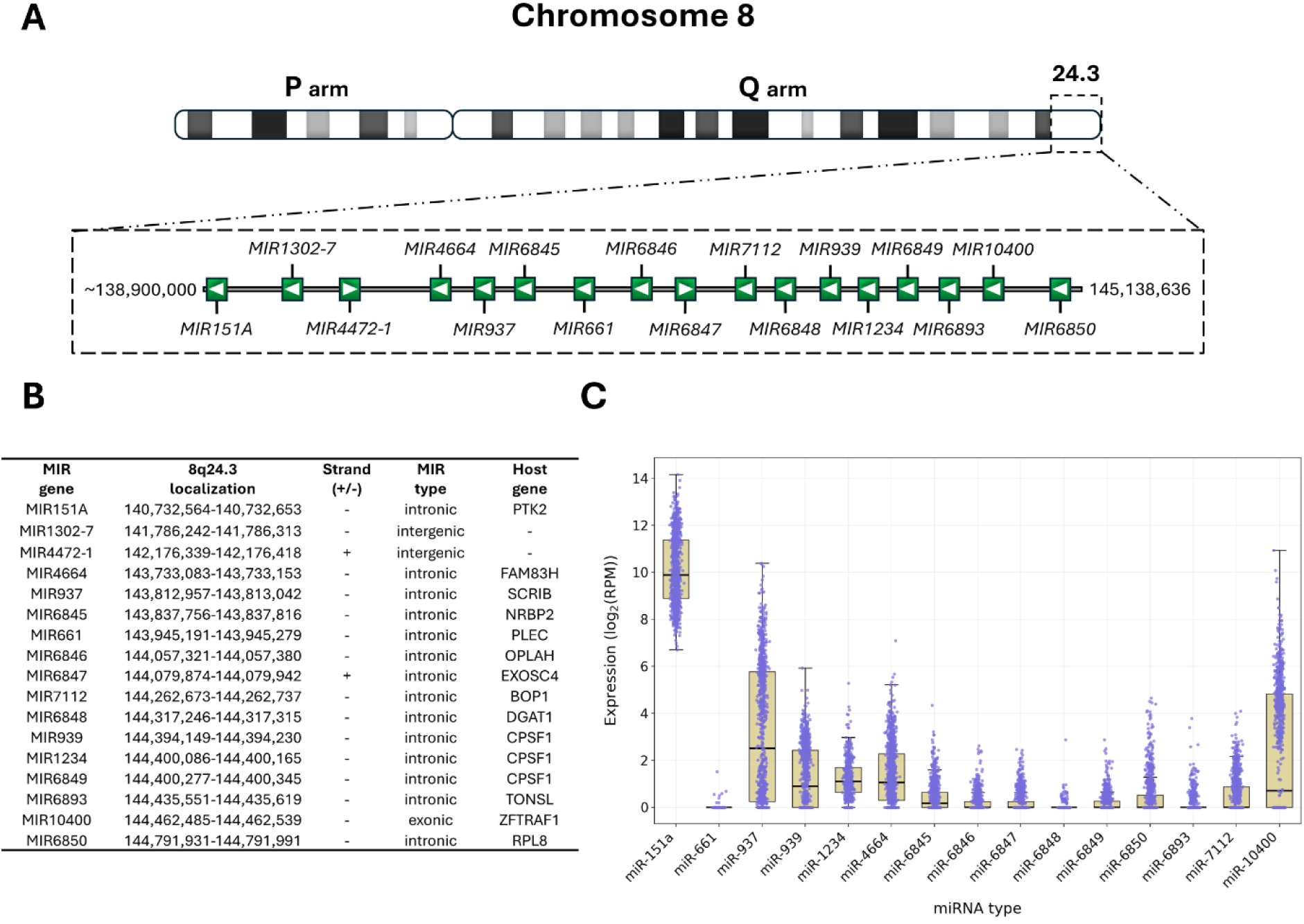
Organization and expression of 8q24.3-encoded miRNAs in ovarian cancer. (A) Schematic map of chromosome 8 and miRNAs genomic positions with strand orientation within the 8q24.3 locus. (B) Annotation table listing 8q24.3-miRNAs with miRNA name, genomic coordinate, strand orientation, miRNA type, and host gene. (C) Box-and-scatter plots showing the distribution of log₂(RPM) expression for 15 miRNAs across TCGA OC tissue samples. Each box represents the interquartile range (IQR) with a black median line; whiskers indicate data within 1.5×IQR. Individual sample values are overlaid as purple dots with slight jitter for visibility, plotted in front of the boxes. miRNAs are ordered numerically on the X axis (lowest to highest).

We obtained miRNA expression data for TCGA HGSOC samples from the DIANA-microRNA Tissue Expression Database (miTED) and noted significant variability among 8q24.3-derived miRNAs (Fig.1C). We found that miR-151a was the most highly expressed miRNA, with median values above 10 log₂(RPM). We also noted that miR-937 had strong expression (median ∼2.5 log₂(RPM)), with a broad range and extreme outliers above 12 log₂(RPM), indicating a lot of variation across patients. miR-939 had an intermediate expression level (median ∼1 log₂(RPM)), while most of the other members of this locus, such as miR-1234, miR-4664, miR-6845, miR-6846, miR-6847, miR-6849, miR-6850, miR-6893, miR-7112, and miR-10400, had low to moderate expression levels, usually below 1 log₂(RPM). In contrast, miR-661 and miR-6848 exhibited consistently low expression levels around background across all tumor samples. These findings indicate that the 8q24.3 locus encodes a substantial array of miRNAs; however, only a specific fraction, including miR-151a, miR-937, and miR-939, has persistently enhanced expression in HGSOC. This indicates that miRNAs within this genomic hotspot may be variably involved in ovarian cancer biology, with certain highly expressed candidates possibly playing functional roles in tumor growth.

To characterize the isoform-specific expression of miRNAs in ovarian cancer tissues, we examined the matched -3p and -5p isoforms of 8q24.3-encoded miRNAs utilizing expression profiles obtained from TCGA (Fig.2). We uncovered extensive asymmetry in strand use, emphasizing the preferential stability and accumulation of certain isoforms among various miRNAs. We observed that miR-151a exhibited strong expression from both strands, with a distinct majority of the -3p isoform, whose median surpassed 12 log₂(RPM), whereas the -5p isoform stayed at 2–3 log₂(RPM). In contrast, miR-937 exhibited significant strand bias, with miR-937-3p attaining a median of 4–5 log₂(RPM) and exceeding 10 log₂(RPM) in certain instances, whilst miR-937-5p was expressed at baseline levels. A reverse trend was seen for miR-939, with the -5p isoform prevailing (median ∼2 log₂(RPM)), whereas the -3p strand was significantly diminished. Likewise, miR-4664 exhibited expression from both strands within the low-to-moderate range (∼1.5–2 log₂(RPM)), but with a little preponderance of the -3p isoform. For miR-6845, we noted an equitable expression of the -3p and -5p isoforms, exhibiting no discernible bias. Numerous miRNAs demonstrated a sustained inclination for the -5p strand, including miR-6846, miR-6847, miR-6848, miR-6849, miR-6850, and miR-6893. In these instances, the -3p isoforms were either significantly diminished (miR-6846 and miR-6847) or almost undetectable (miR-6848, miR-6849, miR-6850, and miR-6893). Both miR-1234 and miR-661 were identifiable solely as singular isoforms. miR-661 exhibited consistently very low expression levels while miR-1234 expression median was approximately ∼1.5 log₂(RPM). Notably, miR-7112 and miR-10400 exhibited significant isoform asymmetry, with expression primarily confined to -3p and -5p strands, respectively.

**Fig. 2.**
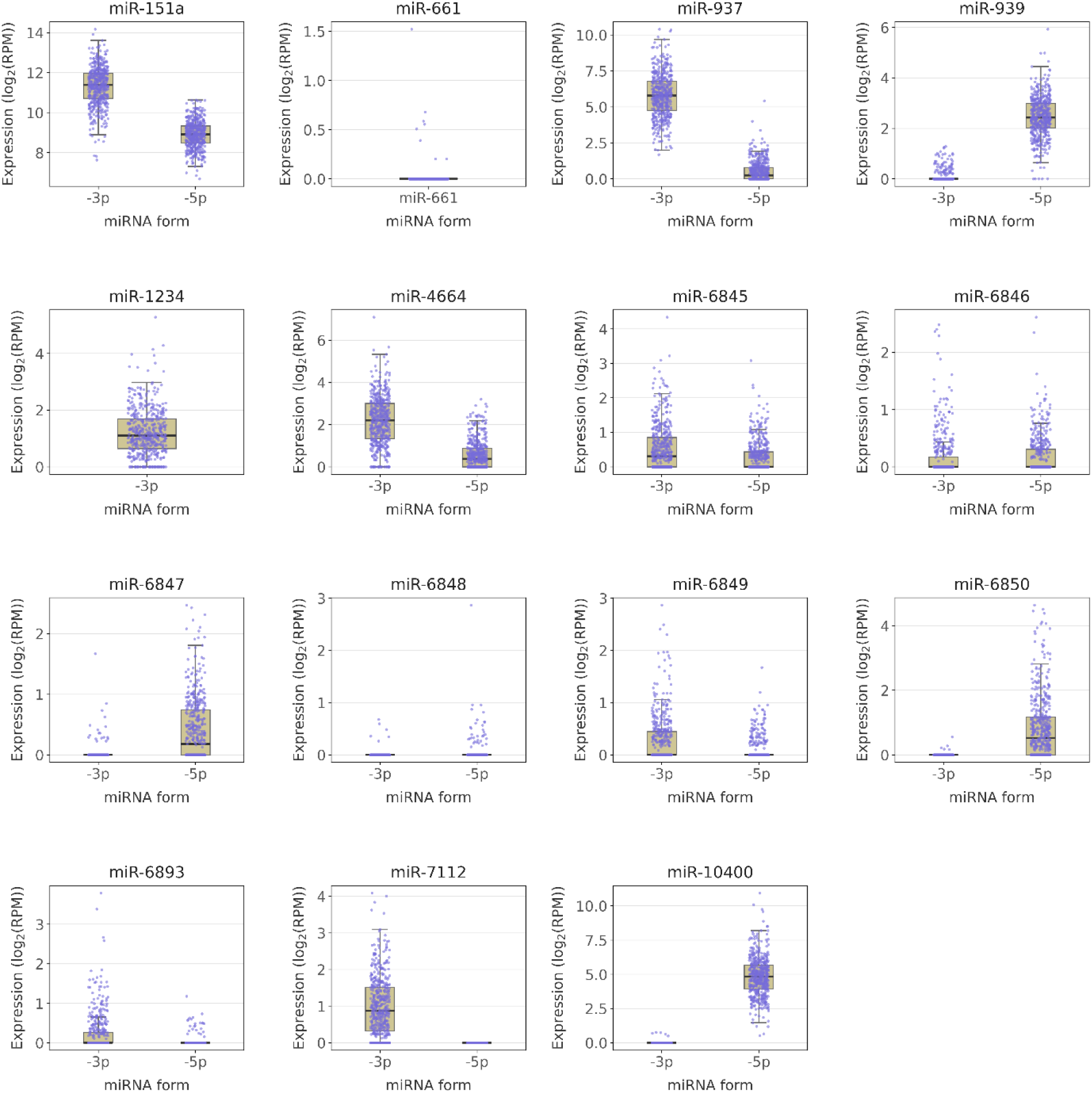
Differential expression of -3p and -5p isoforms across selected miRNAs in OC samples from the TCGA dataset. Box-scatter plot depicts log₂(RPM) expression distribution for both -3p and -5p isoforms of each miRNA in OC samples. For each miRNA (excluding miR-661 and miR-1234), paired isoforms -3p and -5p are shown side-by-side by golden boxes with purple jittered points. Boxes represent the IQR with median lines (black lines); whiskers extend to 1.5× IQR; outliers are hidden for clarity. miRNAs are ordered by numeric ID from lowest to highest.

The extensive inter-sample variability and the existence of upper-tail outliers highlight significant biological heterogeneity among malignancies. The isoform-specific patterns together underscore unique strand selection and stabilization mechanisms, potentially contributing to the functional specialization of 8q24.3-derived miRNAs in the pathophysiology of OC.

Subsequently, we investigated the differential expression of miRNAs encoded by the 8q24.3 locus in ovarian cancer samples that exhibited alterations compared to those without (Suppl. Fig. S2C). As shown in Supplementary Figure S2A, only miR-10400 did not display detectable alterations in the TCGA ovarian cancer cohort; therefore, it was excluded from subsequent analyses. A number of these miRNAs demonstrated markedly elevated expression levels within the altered group. The most significant values were described for miR-939, miR-151A, miR-1234, miR-4664, miR-937, and miR-6850. Less significant, increase was noted for miR-6845, miR-6846, miR-6849, and miR-7112. Additionally, other microRNAs, such as miR-6848, miR-6847, and miR-6893 exhibited significant enrichment in the altered samples. However, it is noteworthy that these findings were accompanied by a greater degree of dispersion and a considerable number of cases with low expression levels. Conversely, miR-151A and miR-937 exhibited elevated expression levels in both altered and unaltered tumor samples. Only miR-661 did not show a statistically significant difference (p = 0.196), maintaining an undetectable level in both altered and unaltered tissues (Suppl. Fig. S2C).

### 3.2 Regulation and functional association of 8q24.3-encoded miRNAs

To refine our understanding of copy number–driven regulation, we next correlated genomic copy number alterations with total miRNA expression using Spearman’s rank correlation coefficient (Fig. 3). This analysis revealed a predominantly positive relationship between copy number gain and miRNA expression, confirming that genomic amplification is a major determinant of transcriptional upregulation within this locus. Across multiple miRNAs, including miR-939, miR-1234, miR-4664, and miR-937, we observed robust positive correlations (0.433 < ρ < 0.546), indicating a strong gene dosage effect. In contrast, isoform-level analysis did not strengthen these associations (Suppl. Fig. S3). The dominant strands showed correlations comparable to, but not exceeding, those observed for total miRNA expression (Suppl. Fig. S3). A second group of miRNAs, such as miR-6850, miR-6845, miR-6846, and miR-6849, exhibited moderate correlations at the total level (0.272 < ρ < 0.350), with isoform-specific analyses yielding similar or slightly weaker associations (Suppl. Fig. S3). Finally, miR-661 (ρ = 0.052) and miR-6848 (ρ = 0.172) displayed weak or negligible correlations, indicating that their expression is likely less dependent on genomic dosage. In several cases, including miR-6845-3p (isoform ρ = 0.290 vs. total ρ = 0.363) and miR-6846-5p (isoform ρ = 0.278 vs. total ρ = 0.318), isoform-specific correlations were modestly reduced (Suppl. Fig. S3), suggesting that strand-level variability may introduce additional noise rather than enhancing sensitivity to copy number changes. Collectively, these findings indicate that copy number amplification exerts a consistent influence on miRNA expression across the 8q24.3 locus, with total miRNA levels providing a robust and stable reflection of genomic dosage. While isoform-specific measurements capture strand-level dynamics, they do not improve and may slightly attenuate the strength of the association with copy number, suggesting that post-transcriptional processes such as strand selection and differential stability introduce variability that is less directly coupled to genomic amplification.

**Fig. 3.**
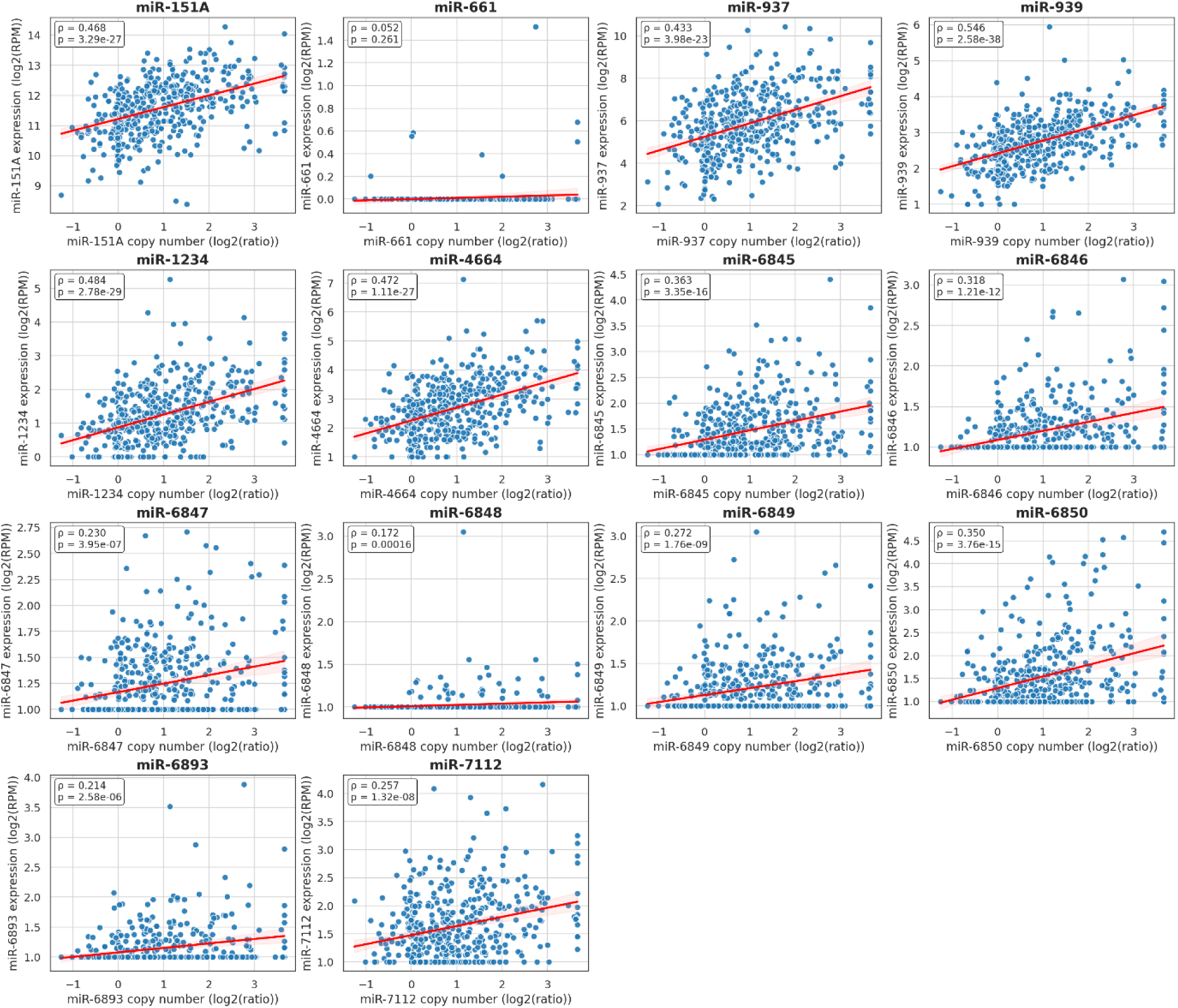
Correlation between copy number variation and 8q24.3-encoded miRNA expression across ovarian cancer samples. Scatter plots show the relationship between copy number (log₂ ratio, X-axis) and expression (log₂ RPM, Y-axis) for individual miRNAs. Each blue dot represents a patient sample, and the red line indicates the linear regression fit with 95% confidence interval shading. Spearman’s correlation coefficient (*ρ*) and corresponding *p*-values are shown in each panel.

To investigate the transcriptional relationships between 8q24.3-encoded miRNAs and their host genes, we examined correlations between total miRNA expression and the corresponding host transcript levels across TCGA OC samples (Fig. 4). This analysis revealed heterogeneous correlation strengths, ranging from absent to strongly positive. Among the analyzed pairs, miR-151A/PTK2 showed the strongest association (ρ = 0.615). Strong positive correlations were also observed for miR-939/CPSF1 (ρ = 0.464) and miR-6846/OPLAH (ρ = 0.422). Moderate correlations were detected for miR-937/SCRIB (ρ = 0.366), miR-1234/CPSF1 (ρ = 0.353), miR-6845/NRBP2 (ρ = 0.319), miR-6893/TONSL (ρ = 0.310), and miR-4664/FAM83H (ρ = 0.310). Weaker positive correlations were found for miR-7112/BOP1 (ρ = 0.221), miR-6850/RPL8 (ρ = 0.216), miR-6848/DGAT1 (ρ = 0.184), and miR-6849/CPSF1 (ρ = 0.171). In contrast, miR-6847/EXOSC4 showed only a weak correlation (ρ = 0.105), while miR-661/PLEC exhibited virtually no association (ρ = 0.028), suggesting that expression of these miRNAs may be relatively uncoupled from host gene transcription.

**Fig. 4.**
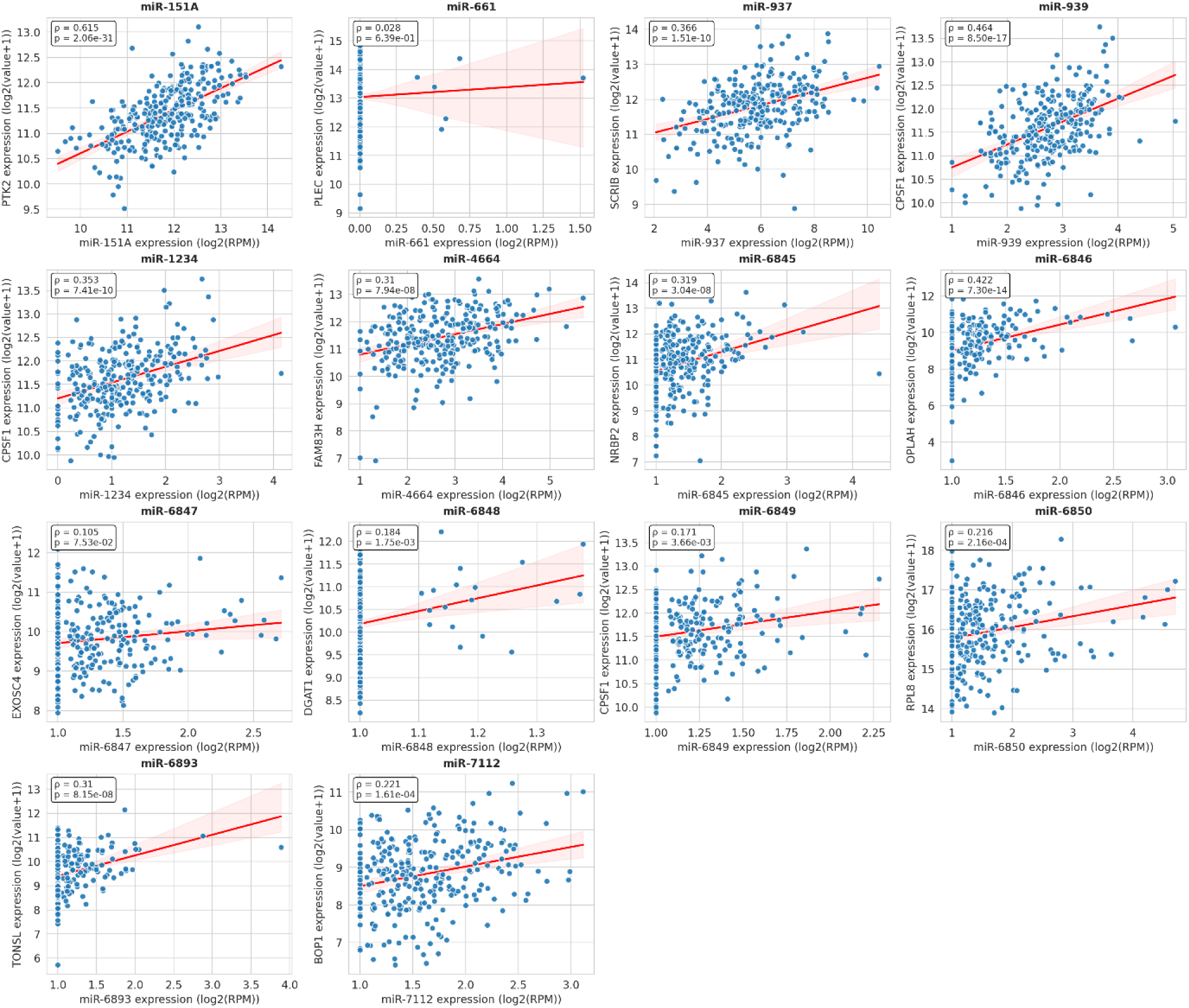
**Expression correlation between 8q24.3-encoded miRNAs and their host gene in ovarian cancer**. Scatterplots showing correlations between the total expression of the 8q24.3-encoded miRNA and its corresponding host gene transcript across TCGA ovarian cancer samples. Each point represents an individual tumor sample; the red line indicates the best-fit linear regression with 95% confidence intervals (light red color). Each panel represents one miRNA–host gene pair. Spearman correlation coefficient (*ρ*) and *p*-value are displayed.

Comparison with the isoform-based analysis showed that, in most cases, the correlation coefficients for the mature isoforms were lower than those obtained using total miRNA expression (Suppl. Fig. S4). This was observed for miR-151A, miR-939, miR-6846, miR-6845, miR-4664, miR-6848, miR-6849, miR-6893, and miR-6850. For miR-937, miR-1234, miR-661, and miR-7112, the correlations were essentially comparable between total miRNA and isoform-level analyses, with only minimal differences. Overall, these findings indicate that total miRNA expression more robustly captures the transcriptional coupling between intronic miRNAs and their host genes, whereas isoform-specific measurements appear to reflect additional post-transcriptional regulatory influences that can attenuate the observed association with host transcript abundance.

Given that MYC is localized at 8q24.21, in close proximity to the 8q24.3 miRNA cluster, we next examined whether expression of these miRNAs was associated with MYC amplification status in ovarian cancer. Overall, the distribution of miRNA expression was shifted toward higher levels in tumors with MYC amplification, indicating that most 8q24.3-encoded miRNAs are more abundant in the amplified subgroup. This pattern was observed for the majority of the analyzed miRNAs, whereas only miR-661 and miR-6848 did not show a significant difference between MYC-amplified and non-amplified samples (Fig. 5). However, this trend was not mirrored by correlation analyses with MYC protein expression, which showed predominantly weak or absent associations and therefore did not support a simple linear relationship between miRNA abundance and MYC protein levels (Suppl. Fig. S5). Together, these results indicate that the higher expression of 8q24.3 miRNAs in MYC-amplified tumors is more consistent with the regional effects of 8q24 copy-number gain than with a direct association with MYC protein output.

**Fig. 5.**
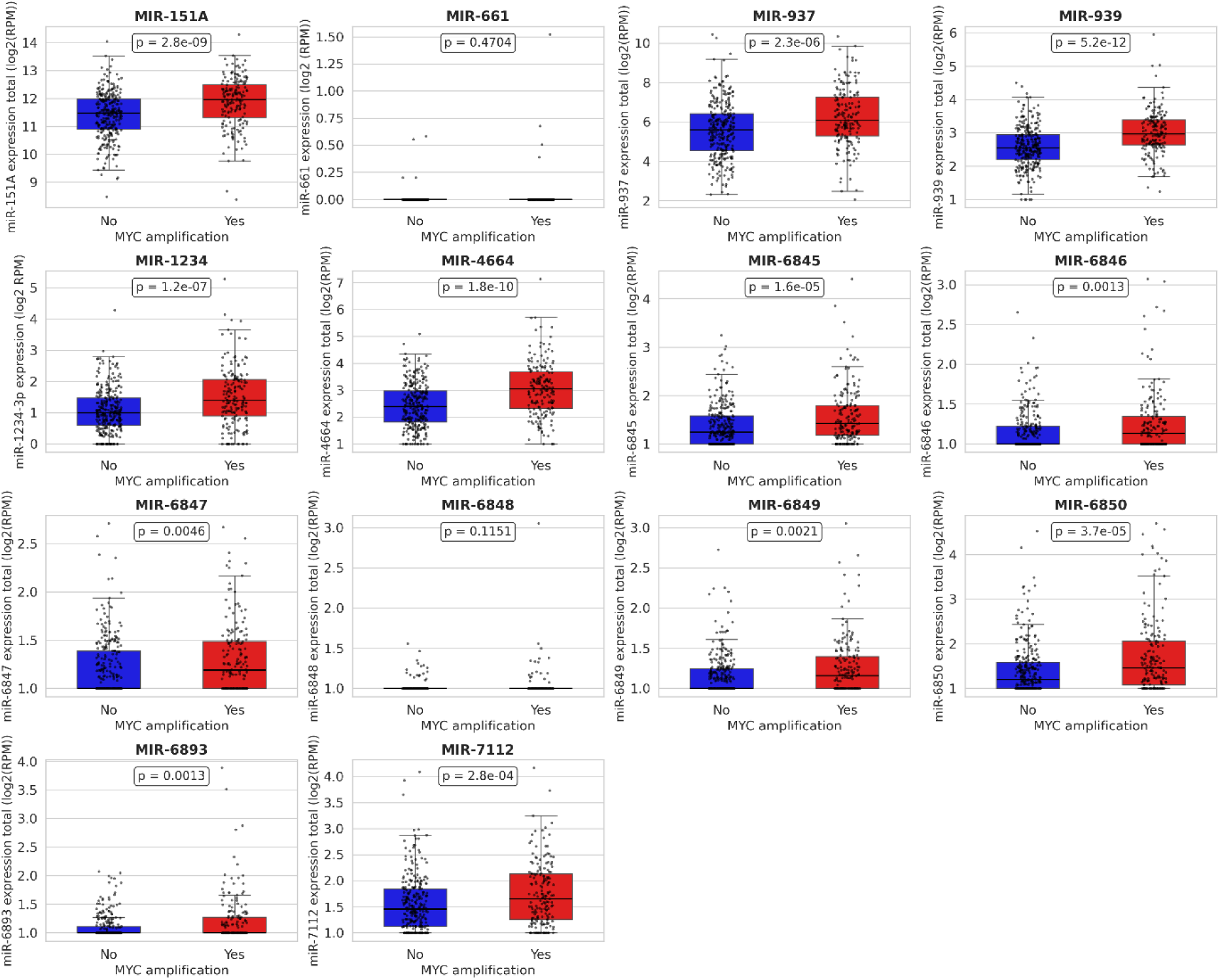
Expression of 8q24.3-encoded miRNAs according to MYC amplification status in ovarian cancer. Boxplots showing the distribution of total expression levels (log2 RPM) of selected 8q24.3-encoded miRNAs in OC samples stratified by MYC amplification status (No vs Yes). Individual data points represent single tumor samples. Boxes indicate the interquartile range, with the median shown as a horizontal line, and whiskers extending to 1.5× the interquartile range. Statistical significance was assessed using the Mann Whitney U test, and *p*-values are indicated for each comparison.

### 3.3 Clinical Associations of 8q24.3-Encoded miRNAs in Ovarian Cancer

To investigate the clinical relevance of 8q24.3-encoded miRNAs in ovarian cancer, we first examined whether their expression differed according to age at diagnosis by stratifying TCGA samples into four groups: <40, 41–60, 61–80, and >81 years (Fig. 5). Overall, most miRNAs showed broadly comparable expression distributions across age categories, although several displayed significant age-related differences. The most evident pattern was observed for miR-151A, whose expression varied significantly across multiple pairwise comparisons and tended to be lower in the oldest group (>81 years) relative to younger and intermediate-age patients. Significant differences were also detected for miR-1234 and miR-4664, both showing differences between the 41–60 and 61–80 groups. In addition, miR-6846 exhibited significant differences involving the >81-year group, indicating that age-related variation is not restricted to a single segment of the cohort. In contrast, the remaining miRNAs did not show statistically significant differences across age groups. Notably, isoform-based analysis also revealed age-associated differences for selected strands, including miR-6849-3p, miR-151A-3p, miR-1234-3p, and miR-4664-3p, largely mirroring the total-miRNA results (Suppl. Table S1). Thus, although age-related variation was not a general feature of the 8q24.3 miRNA locus, a subset of miRNAs showed measurable stratification by patient age at diagnosis.

**Fig. 5.**
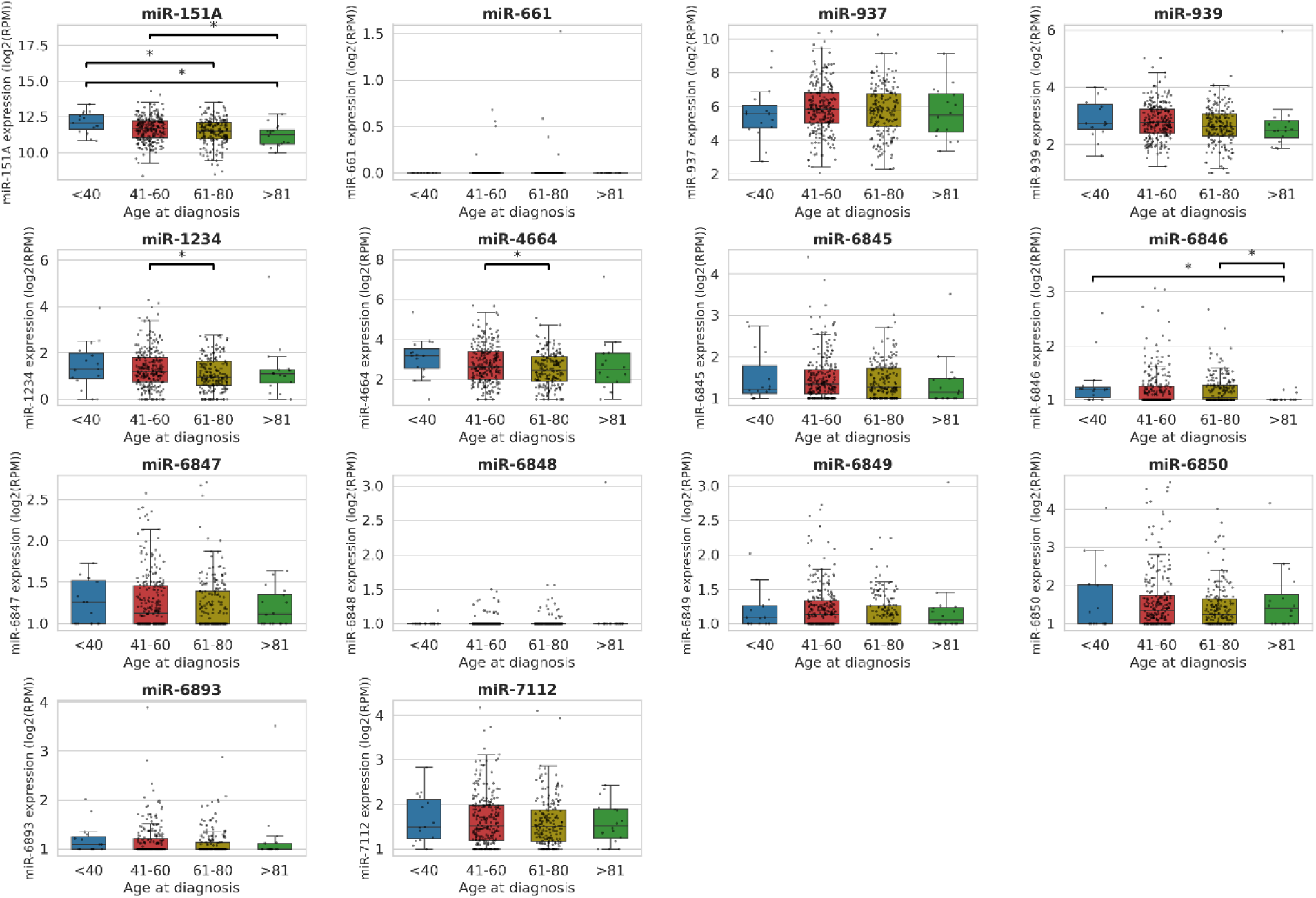
**Expression of 8q24.3-encoded miRNAs in ovarian cancer across diagnosis age groups.** Boxplots showing the distribution of expression levels (log2 RPM) of selected 8q24.3-encoded miRNAs across diagnosis age groups (Groups <40, 41-60, 61-80, >81) in ovarian cancer patients. Individual data points represent single tumor samples. Boxes indicate the interquartile range, with the median shown as a horizontal line, and whiskers extending to 1.5× the interquartile range. Statistically significant differences between diagnosis age groups are indicated (FDR < 0.05).

Next, we assessed the prognostic relevance of 8q24.3-encoded miRNAs by performing Kaplan–Meier analysis of overall survival in patients with HGSOC (Fig. 6). This analysis was conducted using KM Plotter, which does not distinguish between individual miRNA isoforms; therefore, survival associations were evaluated at the level of total miRNA expression. Among the analyzed miRNAs, most did not show a significant association with overall survival. Nevertheless, elevated expression of miR-937, miR-4664, and miR-6849 was associated with a more favorable survival probability, indicating a potential protective prognostic value for these miRNAs in HGSOC. For miR-661 and miR-6893 survival curves suggested a tendency toward improved outcome in the high-expression group, although these associations did not reach statistical significance. By contrast, the remaining miRNAs showed largely overlapping survival distributions between low- and high-expression groups, consistent with the absence of clear prognostic stratification. Notably, none of the analyzed miRNAs displayed a significant association with poorer survival when highly expressed. Collectively, these findings identify miR-937, miR-4664, and miR-6849 as the most promising candidates for favorable prognostic biomarkers among the 8q24.3-encoded miRNAs examined in HGSOC.

**Fig. 6.**
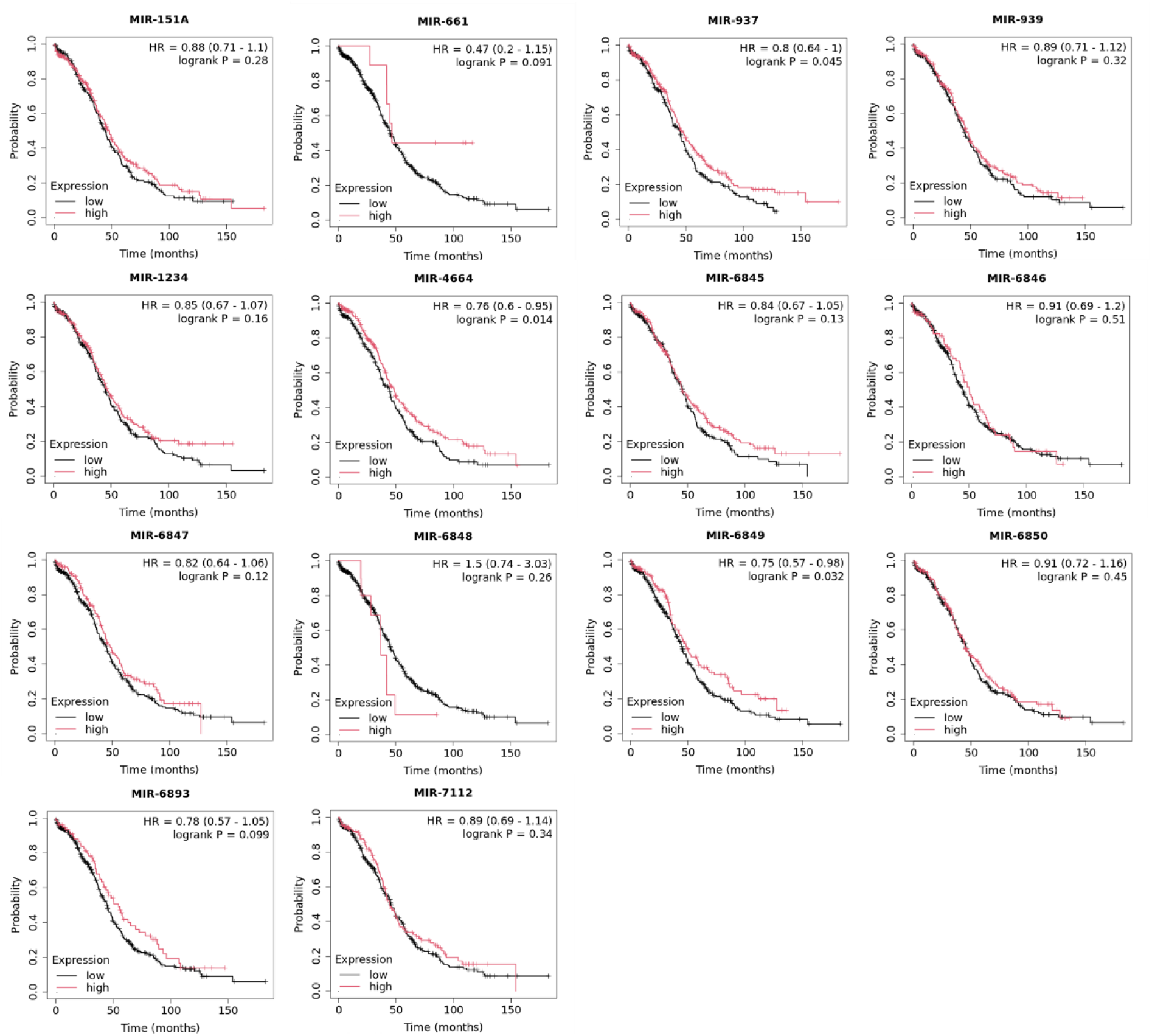
Kaplan–Meier overall survival curves for ovarian cancer patients stratified by expression of 8q24.3-encoded miRNAs. HGSOC patients were divided into high- and low-expression groups (red = high, black = low) for each miRNA, and overall survival was compared using log-rank tests. Hazard ratios (HR) with 95% confidence intervals and *p*-values are shown for each plot.

To assess whether 8q24.3-encoded miRNAs are differentially expressed across EOC histotypes, we compared total miRNA expression among the four major subtypes: endometrioid, serous, mucinous, and clear cell carcinoma (Suppl. Fig. S6). Overall, most miRNAs showed broadly overlapping expression distributions across subtypes, indicating that expression of the 8q24.3 miRNA locus is largely conserved among EOC histotypes. No clear subtype-restricted cluster of highly expressed miRNAs emerged at the total miRNA level, although a tendency toward slightly higher expression in the endometrioid group could be observed for miR-937, miR-939, and miR-6893, whereas miR-151A, miR-6846, and miR-7112 showed somewhat higher median expression in the mucinous subtype. In contrast, the serous subtype often displayed intermediate-to-lower expression, while clear cell samples were generally more variable, likely reflecting the smaller sample size in this group. After statistical refinement, only miR-7112 retained a significant subtype-associated difference, distinguishing mucinous from serous tumors (Suppl. Fig. S6 and Suppl. Table S2). All other miRNAs showed no significant differences among the analyzed histotypes. Importantly, isoform-resolved analysis likewise did not identify any significant subtype-associated miRNA candidate, indicating that strand-level evaluation did not improve discrimination among EOC subtypes in this dataset (Suppl. Table S2). Thus, both total miRNA and isoform-specific analyses indicate that 8q24.3-encoded miRNAs have only limited utility for distinguishing EOC histotypes, with miR-7112 representing the only miRNA showing a detectable subtype-related difference at the total expression level. Finally, we examined the association between 8q24.3-encoded miRNA expression and additional clinicopathological variables, including FIGO (International Federation of Gynecology and Obstetrics) stage, patient ethnicity, tumor residual disease, lymphatic invasion, mean tumor dimension, and chemotherapy exposure, to determine whether these miRNAs are linked to clinical features beyond subtype, age at diagnosis, and survival. Overall, these analyses revealed only limited associations, with substantial overlap in miRNA expression distributions across the compared groups. No significant associations were observed for FIGO stage (Suppl. Table S3), patient ethnicity (Suppl. Table S4), or tumor residual disease (Suppl. Table S5), indicating that expression of 8q24.3-encoded miRNAs is largely independent of these parameters in this cohort. Likewise, chemotherapy exposure did not show any significant relationship with miRNA expression. Chemotherapy was common in this cohort, and most treated patients received standard platinum–taxane-based regimens; however, no statistically significant differences in miRNA levels were detected across treatment groups, suggesting that treatment exposure had little measurable impact on expression of this miRNA set. Among the additional variables examined, only lymphatic invasion (Suppl. Table S6) and mean tumor dimension (Suppl. Table S7) showed significant associations, and these were restricted to miR-151A. Specifically, both total miR-151A expression and miR-151A-3p expression differed significantly according to lymphatic invasion status. In addition, total miR-151A expression and miR-151A-3p showed weak but significant positive correlations with mean tumor dimension, indicating that higher expression of this miRNA is modestly associated with larger tumor size. Taken together, these findings indicate that most of the additional clinicopathological variables examined, including FIGO stage, ethnicity, tumor residual disease, and chemotherapy exposure, exert limited influence on the expression of 8q24.3-encoded miRNAs. The only consistent signal was observed for miR-151A, suggesting that this miRNA may be more closely linked to selected features of tumor progression than the other members of the 8q24.3 miRNA set.

### 3.4 Functional enrichment analysis of 8q24.3-encoded miRNA target genes

To gain functional insight into the biological roles of 8q24.3-encoded miRNAs, we performed over-representation analysis on the experimentally validated target genes from miRTarBase v10.0. This analysis was done to uncover whether these miRNAs’ target genes converge on specific biological processes and/or pathways relevant to ovarian cancer.

The GO Biological Processes enrichment analysis demonstrated that the investigated miRNAs were substantially implicated in cellular stress responses and cell fate regulation (FDR<0.05) (Fig. 8A). Among them were *DNA integrity checkpoint signaling, cellular senescence*, and *regulation of proteolysis involved in protein catabolic processes.* This suggests that these 8q24.3-encoded miRNAs may play a role in controlling genomic stability and proteostasis. Further enrichment of these miRNAs pointed to altered *innate immune and signaling responses*, including the *cytoplasmic pattern recognition receptor signaling pathway* and *intracellular receptor signaling pathway*, suggesting that 8q24.3 miRNAs may fine-tune inflammatory signaling and ligand-responsive transcriptional programs in ovarian cancer cells. Enrichment of processes related to circadian regulation, such as *entrainment of the circadian clock by photoperiod* and *photoperiodism*, further implicates these miRNAs in the deregulation of temporal control of cell cycle progression, metabolism, and DNA repair. Metabolic adaptation was also reflected by enrichment of the *serine family amino acid biosynthetic process*, a pathway closely linked to nucleotide synthesis and redox balance in cancer cells [33].

**Fig. 8.**
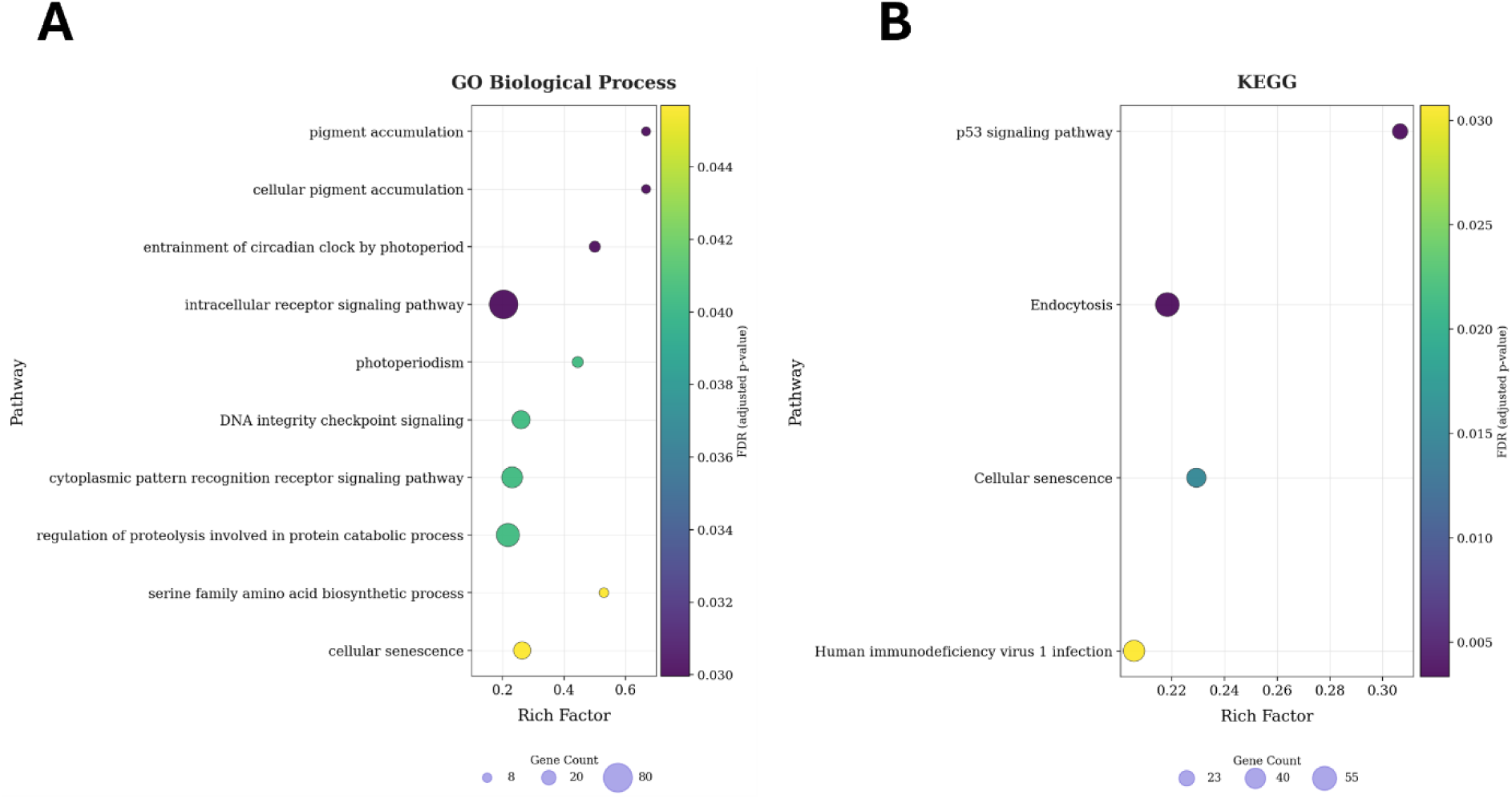
Gene Ontology and KEGG enrichment analysis of 8q24.3-encoded miRNAs. Bubble plots display the top enriched GO Biological Process terms (A) and KEGG pathways (B) identified by ORA. Bubble size represents the number of enriched genes (gene count), while bubble color corresponds to the FDR-adjusted p-value. Only significantly enriched terms (FDR-adjusted p<0.05) are shown on the graphs.

Consistently, KEGG pathway enrichment analysis indicated that significantly enriched pathways for 8q24.3-encoded miRNA genes were *endocytosis*, *p53 signaling pathway*, and *cellular senescence* (Fig. 8B). Given the disruption of p53 signaling in HGSOC, enrichment of this pathway underscores the involvement of 8q24.3 miRNAs in regulating downstream components of genome stability, stress responses, and cell fate decisions. Enrichment of endocytosis suggests additional roles in intracellular trafficking and signal modulation, processes critical for tumor cell adaptation and survival.

In the Hallmark pathway analysis, the most represented were *TNF-α signaling via NF-κB*, *estrogen response*, *mTORC1 signaling*, *p53 pathway*, *PI3K/Akt/mTOR pathway*, and *G2/M checkpoint* (Suppl. Fig. S7). However, unlike the GO and KEGG results, none of the Hallmark pathways reached the statistical significance (FDR > 0.05).

Taken together, these results suggest that 8q24.3-encoded miRNAs might modulate tumor cell plasticity and survival under stress conditions of chronic genomic and metabolic stress.

## 4. Discussion

Ovarian cancer remains one of the most lethal malignancies in women, largely because most patients are diagnosed at advanced stages and because HGSOC is characterized by marked genomic instability [5,31]. In this context, recurrent gain of chromosome 8 and, in particular, amplification of the 8q24.3 cytoband represent prominent genomic features of HGSOC [32]. However, despite increasing interest in this region, its non-coding RNA output has remained insufficiently characterized. In the present study, we performed an integrative analysis of all miRNAs encoded at 8q24.3 and show that this locus does not behave as a uniformly activated amplicon. Rather, it constitutes a heterogeneous regulatory compartment in which genomic dosage, host-gene context, and post-transcriptional processing collectively shape mature miRNA abundance.

A first important finding of our study is that the 8q24.3 locus contains a sizeable and structurally diverse repertoire of miRNAs, but only a subset is consistently abundant in HGSOC. Among the analyzed candidates, miR-151A, miR-937, and miR-939 showed the highest expression, whereas several other miRNAs were present at low or near-background levels despite residing within the same amplified region. This observation indicates that copy-number gain alone is not sufficient to explain mature miRNA output. Such heterogeneity is consistent with the broader literature showing that cancer-associated miRNA deregulation reflects multiple layers of control, including transcriptional regulation [13], epigenetic remodeling [16], altered processing by the miRNA biogenesis machinery [17], and contextual influences from the tumor microenvironment [34]. Thus, our data refine a simple gene-dosage model and support the view that 8q24.3 amplification creates a permissive genomic context, but that individual miRNAs remain subject to additional regulatory constraints.

Our isoform-resolved analyses further underscore this regulatory complexity. We observed marked asymmetry in strand usage across several 8q24.3-encoded miRNAs, with some precursors producing a strongly dominant mature strand and others showing more balanced processing. These patterns support the idea that strand selection and strand stability are important determinants of mature miRNA accumulation in ovarian cancer, in line with current concepts of arm selection and isomiR regulation [35,36]. At the same time, the isoform-level quantification did not strengthen the association between copy number and expression. On the contrary, total miRNA measurements generally provided correlations that were equal to or stronger than those observed for individual dominant strands. A similar pattern emerged in the host-gene analysis, where total miRNA expression more robustly captured transcriptional coupling with host transcripts than isoform-specific measurements. Taken together, these findings suggest that isoform-resolved profiling is highly informative for understanding post-transcriptional diversification, but not necessarily superior for detecting dosage-dependent regulation. In this setting, strand-specific measurements appear to capture additional biological variability that is only partially linked to copy-number gain.

The transcriptional relationships between intronic miRNAs and their host genes also revealed substantial heterogeneity within the locus. While some pairs, such as miR-151A/PTK2, miR-939/CPSF1, and miR-6846/OPLAH, showed moderate to strong positive correlations, other miRNA-host gene pairs were only weakly coupled or essentially uncoupled. These data argue against a single transcriptional logic across the 8q24.3 region and instead support the existence of distinct regulatory architectures among neighboring miRNAs. Such variability may reflect variable chromatin accessibility [13], the presence of independent promoters [37], integrity of the precursor RNA stem [37], or differential methylation. Indeed, previous studies have shown that amplified regions in HGSOC often undergo epigenetic remodeling, including CpG methylation that can silence or attenuate miRNAs despite high gene dosage [38]. This interpretation is reinforced by our observation that miRNA abundance was broadly higher in MYC-amplified tumors, whereas correlations with MYC protein expression were weak or absent. Therefore, the association with MYC amplification is more plausibly explained by regional 8q gain than by a direct linear relationship between MYC protein levels and miRNA output. These findings underscore that even within a dense cluster, individual miRNAs can be controlled by distinct regulatory programs.

The clinical analyses further indicate that the 8q24.3 miRNA cluster has selective, rather than widespread, clinicopathological relevance. Most miRNAs showed no significant associations with FIGO stage, ethnicity, tumor residual disease, or chemotherapy exposure, suggesting that expression of this locus is largely independent of these variables in the analyzed cohort. Likewise, histotype-associated differences were limited. At the total miRNA level, only miR-7112 retained a significant difference among EOC histotypes, and isoform-resolved analysis did not improve subtype discrimination. These results suggest that, at least in currently available datasets, 8q24.3-derived miRNAs have limited utility as broad histotype classifiers. Age-at-diagnosis analyses revealed only limited associations involving a subset of miRNAs, including miR-151A, miR-1234, miR-4664, and miR-6849. Notably, isoform-based analysis identified significant age-related differences for miR-151A-3p, miR-1234-3p, miR-4664-3p, and miR-6849-3p, broadly paralleling the total-miRNA results and suggesting that these associations are not solely driven by aggregate expression. However, these findings should be interpreted cautiously, as the youngest (<40 years) and oldest (>81 years) patient groups were relatively small, likely reducing statistical power and increasing variability at the extremes.

Clinical stratification analyses further contextualize these molecular findings. Associations between miRNA expression and patient age at diagnosis or ethnicity suggest that regulatory variation at the 8q24.3 locus may intersect with demographic and biological heterogeneity in ovarian cancer. Previous epidemiological studies have reported differences in ovarian cancer incidence, age of onset, and outcome across racial groups [39]. While our observations should be interpreted cautiously due to cohort composition and sample size limitations, they underscore the importance of integrating clinical covariates when evaluating miRNA-based biomarkers and caution against assuming universal applicability across patient populations.

Among the clinicopathological variables examined, miR-151A was the only miRNA to show a more consistent association with parameters linked to tumor progression, including lymphatic invasion and mean tumor dimension. Although these correlations were weak, they raise the possibility that miR-151A may be more closely linked to local tumor behavior than other members of the locus. This is notable because miR-151A was also the most abundant 8q24.3-encoded miRNA in TCGA dataset, suggesting that both expression level and regulatory integration may contribute to its biological relevance. Nevertheless, the absence of broader associations across the remaining clinicopathological variables argues against a generalized clinical impact of all 8q24.3-resident miRNAs and instead supports a model in which only selected members acquire context-dependent relevance.

Our survival analysis identified miR-937, miR-4664, and miR-6849 as candidate favorable prognostic markers, since higher total expression of these miRNAs was associated with improved overall survival in HGSOC[2]; however, overall outcomes remain extremely poor because most patients are diagnosed at an advanced stage of disease. Moreover, Kaplan-Meier analyses were based on total miRNA measurements and therefore could not distinguish potentially divergent contributions of individual strands. This issue may be particularly relevant for miR-937. While our data associate higher total miR-937 expression with better survival, Zhang et al. recently reported that MIR937 amplification can promote ovarian cancer progression through autophagy-related mechanisms 𝔆𝔆. One possible explanation for this apparent discrepancy is that total-expression measurements are dominated by the more abundant strand, whereas the biologically active isoform in a given context may differ. More broadly, these observations are consistent with the literature indicating that miRNAs can exert tumor-suppressive or tumor-promoting functions depending on tissue context, target availability, and regulatory state𝔆𝔆. Thus, the prognostic signal observed here should be regarded as hypothesis-generating rather than definitive evidence of a uniformly protective function.

The functional enrichment analysis provides a plausible biological framework for these observations. Experimentally validated targets of 8q24.3-encoded miRNAs converged on pathways related to stress responses, senescence, p53 signaling, endocytosis, and metabolic adaptation. These processes are highly relevant to HGSOC biology, where chronic replication stress, defective p53 signaling, and adaptive rewiring of survival pathways are central features [3–5]. Notably, although Hallmark categories such as PI3K/AKT/mTOR, p53 pathway, and G2/M checkpoint did not reach statistical significance after correction, they showed coherent trends that were broadly consistent with the significant GO and KEGG results. This pathway-level convergence also fits with the established role of tumor-suppressive miRNAs as network regulators that restrain oncogenic signaling, stress adaptation, and tumor-promoting cell-state transitions [40,41]. In this regard, our previous functional study on miR-6850 is particularly informative, as ectopic miR-6850 expression in HGSOC models reduced proliferation, perturbed cell-cycle progression, and suppressed PI3K/AKT/mTOR signaling [22]. Although miR-6850 was not associated with survival in the present cohort, these experimental data support the idea that at least some members of the 8q24.3 cluster can exert tumor-restraining effects despite residing within an amplified chromosomal region.

This study has several limitations. First, it is based on retrospective in silico analyses of bulk tumor datasets and therefore does not permit causal inference. Second, the use of total miRNA measurements in some clinical datasets, including survival analysis, limits the ability to resolve strand-specific functions. Third, the multicopy nature of some loci prevented locus-specific interpretation for selected miRNAs. Fourth, bulk transcriptomic data cannot fully capture the contribution of stromal and immune compartments, even though miRNA deregulation is increasingly recognized as being shaped by tumor-microenvironmental interactions [34]. Finally, the observed associations require direct wet-lab validation and mechanistic follow-up. Future studies should therefore combine locus-specific and isoform-specific quantification with functional perturbation, epigenetic profiling, and microenvironment-aware experimental models to determine which 8q24.3-encoded miRNAs are true drivers, biomarkers, or bystanders in ovarian cancer.

## 5. Conclusions and perspectives

This study provides the first comprehensive analysis of all miRNAs encoded within the 8q24.3 locus in ovarian cancer and shows that this region functions as a heterogeneous non-coding regulatory hub rather than a passive consequence of chromosomal amplification. By integrating copy-number, expression, host-gene, histotype, and survival analyses, we show that miRNA output from 8q24.3 is shaped by both genomic dosage and post-transcriptional regulation. Although isoform-level analyses revealed marked strand asymmetry and regulatory complexity, total miRNA expression more robustly reflected copy-number and host-gene relationships. Among the analyzed candidates, miR-937, miR-4664, and miR-6849 emerged as promising but still preliminary prognostic markers in HGSOC. Overall, these findings expand the functional landscape of the 8q24.3 locus and provide a basis for future validation and mechanistic studies of its miRNAs in ovarian cancer.

## Author Contributions

All authors contributed to the conceptualization and research design. KF and MP designed the research process. KF, IM, and FC performed the analysis. KF, IM, and FC analyzed the data. KF and MP wrote the original draft, reviewed, and edited the paper. KF, IM, FC, and MP read and approved the final version of the submitted manuscript.

## Acknowledgments

The authors gratefully acknowledge the networking efforts of the TRANSLACORE Action (CA21154) by COST (European Cooperation in Science and Technology), which facilitated their professional connection.

## Availability of data

The data used to support the findings of this scientific work are available from the corresponding author upon reasonable request.

