## Supplementary figures for "“Landscape of 8q24.3-Encoded microRNAs and Their Prognostic Impact in Ovarian Cancer”"

Supplementary data:

**Figure S1**

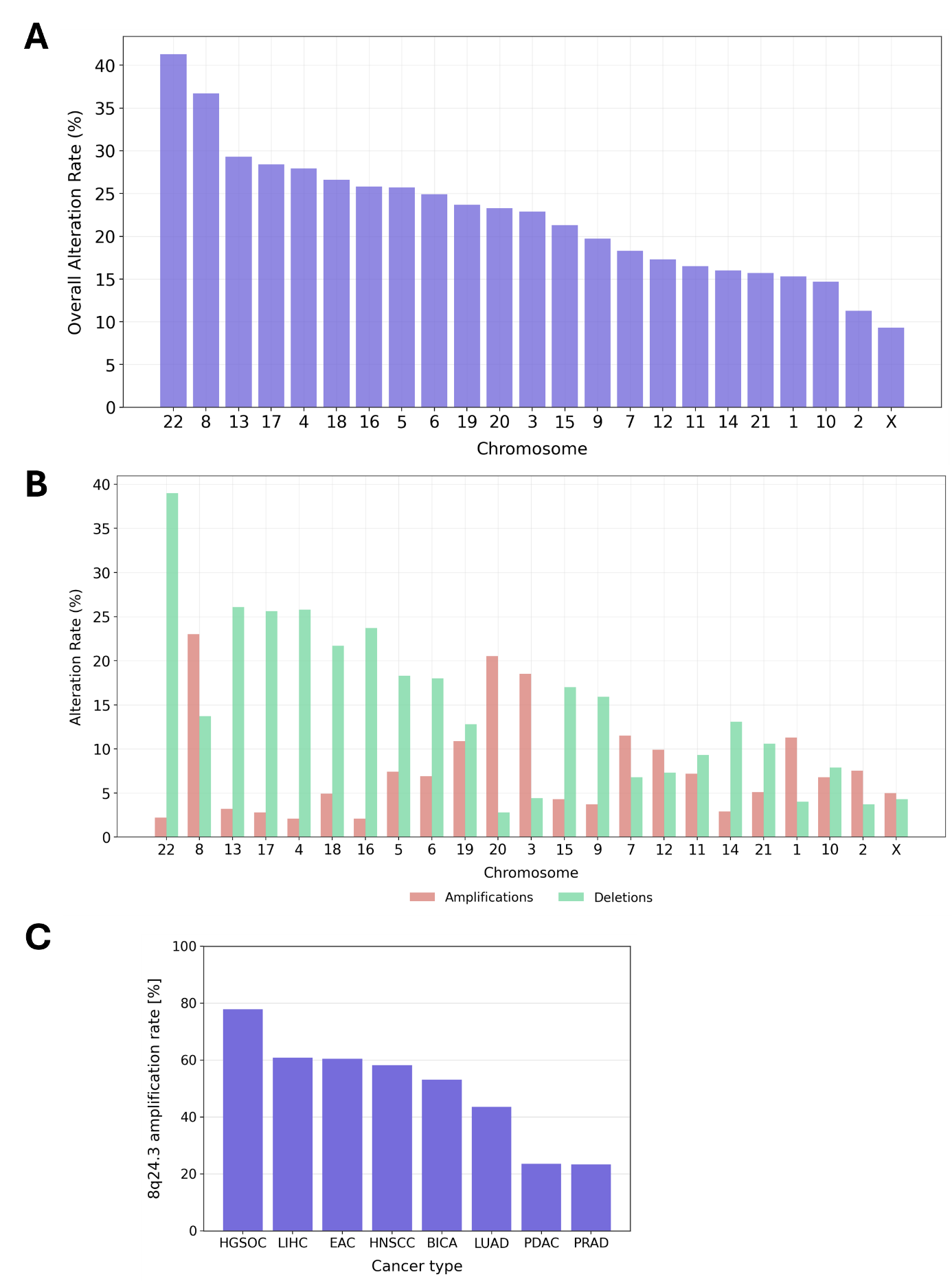

**Fig.S1 Landscape of chromosomal alterations in OC samples from TCGA dataset**. (A) Overall alteration rates across human chromosomes. Chromosomes were sorted out from the highest to the lowest overall alteration rate. (B) Distribution of alteration types (amplification vs deletions) across human chromosomes. The amplification bars are marked with red color while deletion bars are marked with green color. (C) Genetic amplification of 8q24.3 locus across human cancers retrieved from TCGA PanCancer atlas. HGSOC – High Grade Serous Ovarian Cancer; LIHC – Liver Hepatocellular Carcinoma; EAC – Esophageal Adenocarcinoma; HNSCC – Head and Neck Squamous Cell Carcinoma; BICA – Breast Invasive Carcinoma; LUAD – Lung Adenocarcinoma; PDAC – Pancreatic Adenocarcinoma; PRAD – Prostate Adenocarcinoma

**Figure S2**

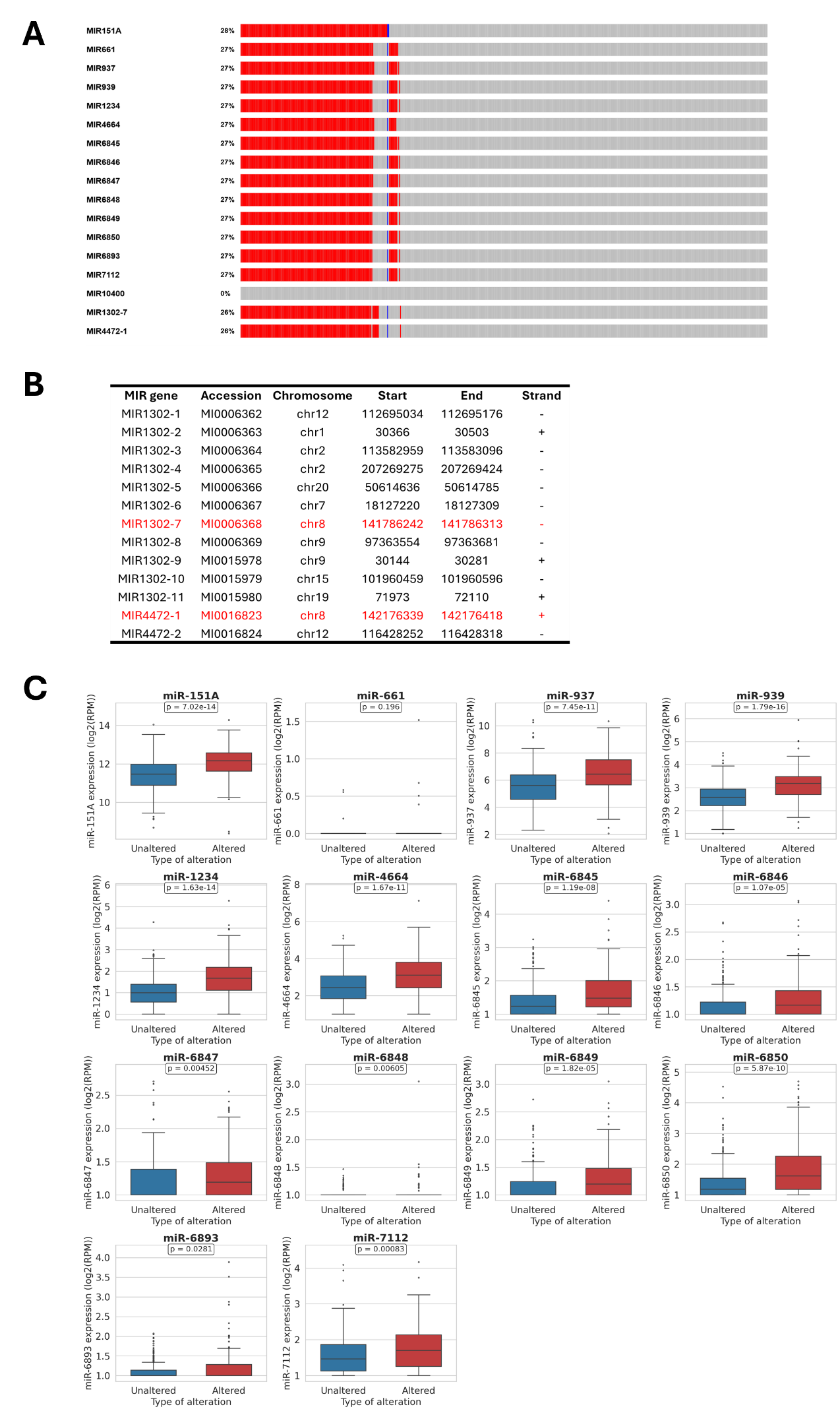

**Fig.S2 Genetic alternation and expression of selected 8q24.3 miRNAs in OC samples from TCGA dataset.** (A) Genetic alternation profiles of 8q24.3-encoded miRNAs in TCGA OC samples. (B) Annotation table of miR-1302 and miR-4472 family within human genome. Table shows miRNA gene name, miRBase accession number, chromosomal localization, start and end of miRNA, and strand orientation. MiR-1302-7 and miR-4472-1 are highlighted in red as they are localized on 8q24.3 chromosome. (C) Comparison of miRNAs expression in altered and unaltered samples. Boxplots represent log₂ RPM expression levels for individual miRNAs in ovarian cancer tissues classified as altered or unaltered. Each subplot corresponds to a single miRNA, with the central line indicating the median expression, boxes representing the IQR, and whiskers extending to 1.5× IQR. Statistical significance was assessed using the Mann Whitney U test, and p-values are indicated for each comparison.

**Figure S3**

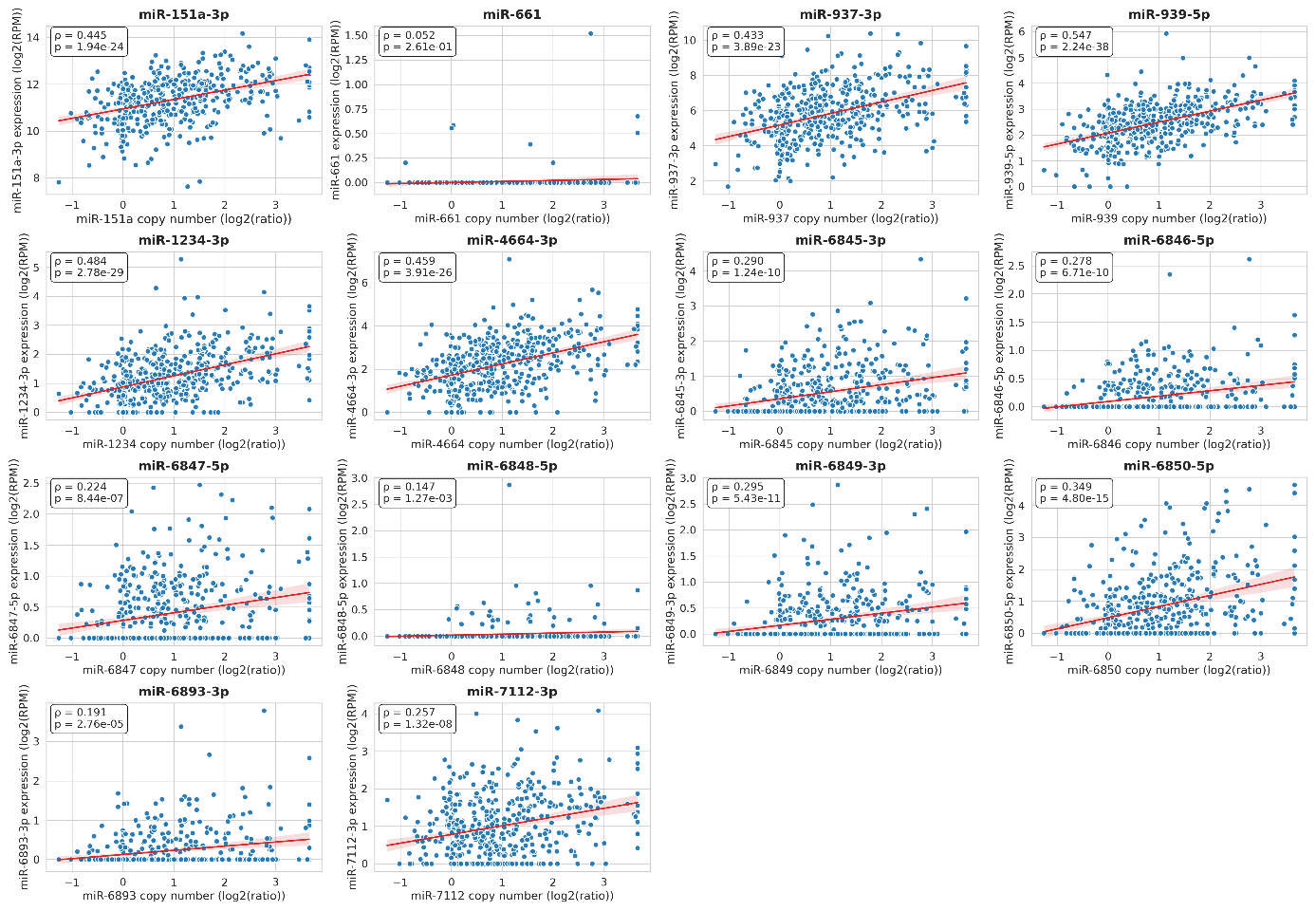

**Fig. S3 Isoform-specific correlation between 8q24.3-encoded miRNA copy number variation and their expression in OC.** Scatter plots show the relationship between copy number (log₂ ratio, X-axis) and expression (log₂ RPM, Y-axis) for individual miRNAs. Each blue dot represents a patient sample, and the red line indicates the linear regression fit with 95% confidence interval shading. Spearman’s correlation coefficient (ρ) and corresponding p-value are shown in each panel.

**Figure S4**

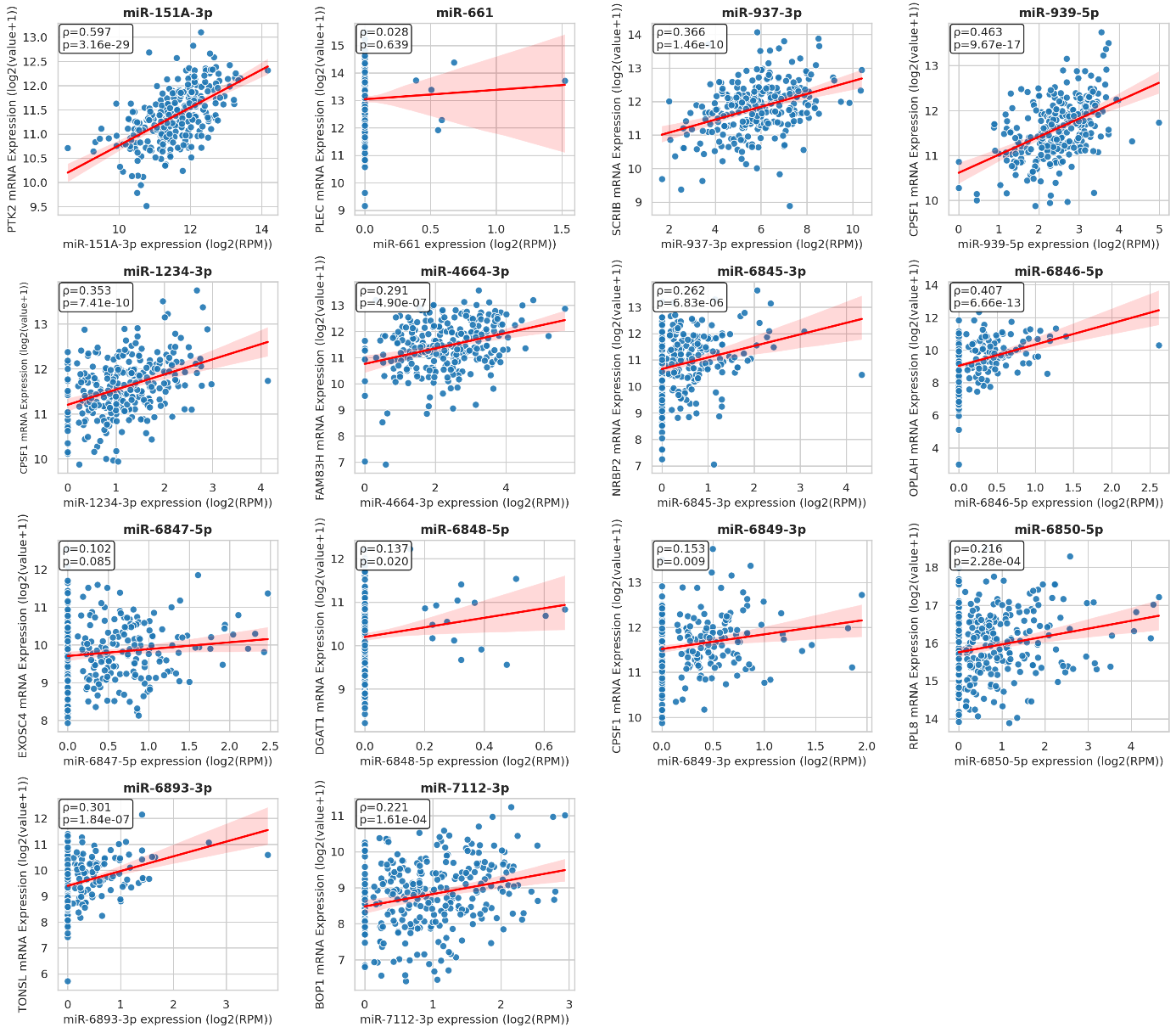

**Fig. S4 Isoform-specific correlation between 8q24.3-encoded miRNAs and their host gene expression in HGSOC**. Scatterplots depict correlations between the expression levels of selected miRNAs located within the 8q24.3 locus and their respective host genes in HGSOC samples from TCGA. miRNA expression (log₂ RPM, X-axis) is plotted against host gene mRNA expression (log₂[Value+1], Y-axis). Each panel represents one miRNA–host gene pair with the Spearman correlation coefficient (ρ) and *p*-value indicated.

**Figure S5**

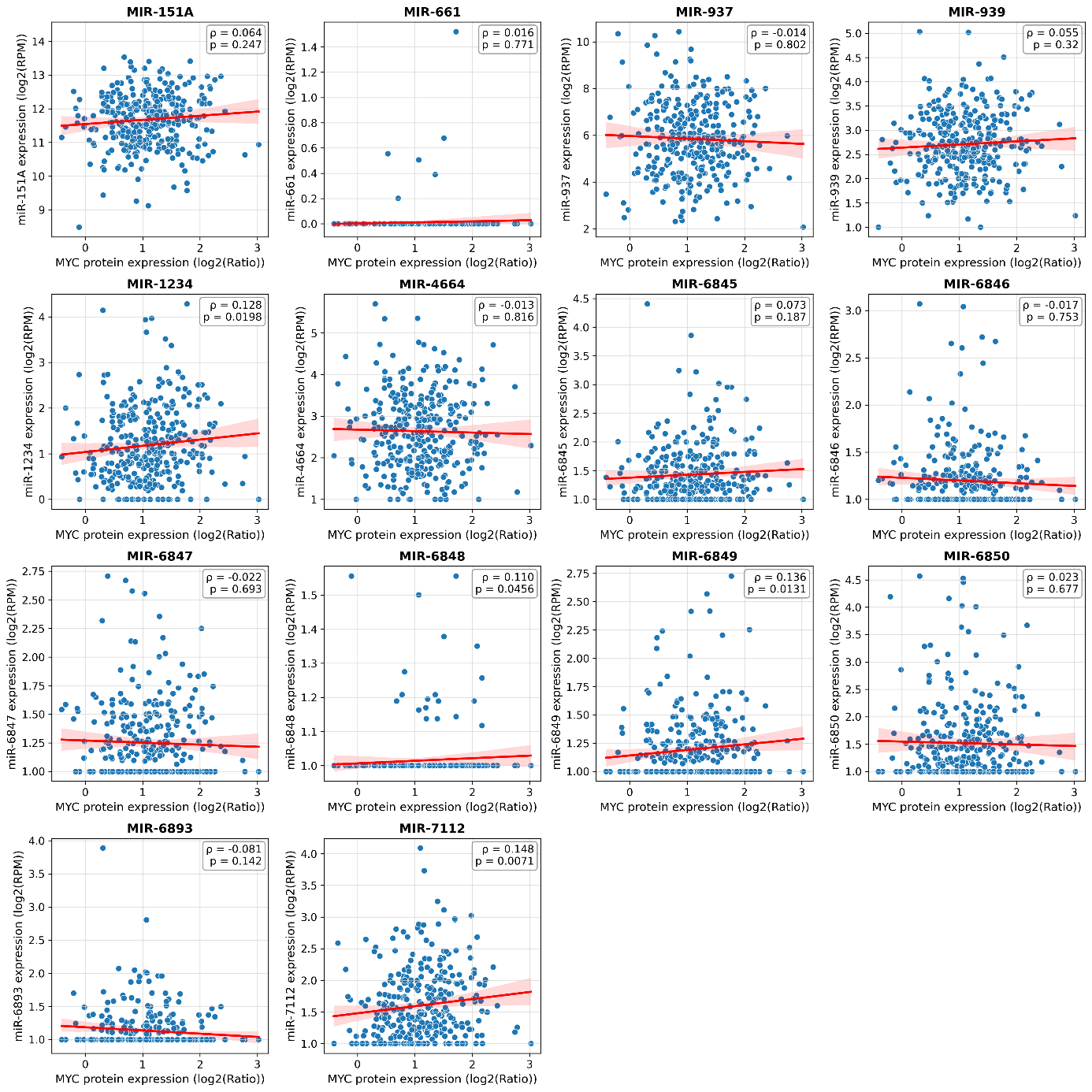

**Fig. S5 Expression correlation between 8q24.3-encoded miRNAs and MYC protein in ovarian cancer**. Scatterplots showing correlations between the expression of the 8q24.3-encoded miRNA and MYC protein across TCGA ovarian cancer samples. Each point represents an individual tumor sample; the red line indicates the best-fit linear regression with 95% confidence intervals (light red color). Each panel represents one miRNA–MYC pair. Spearman correlation coefficient (ρ) and *p*-value are displayed.

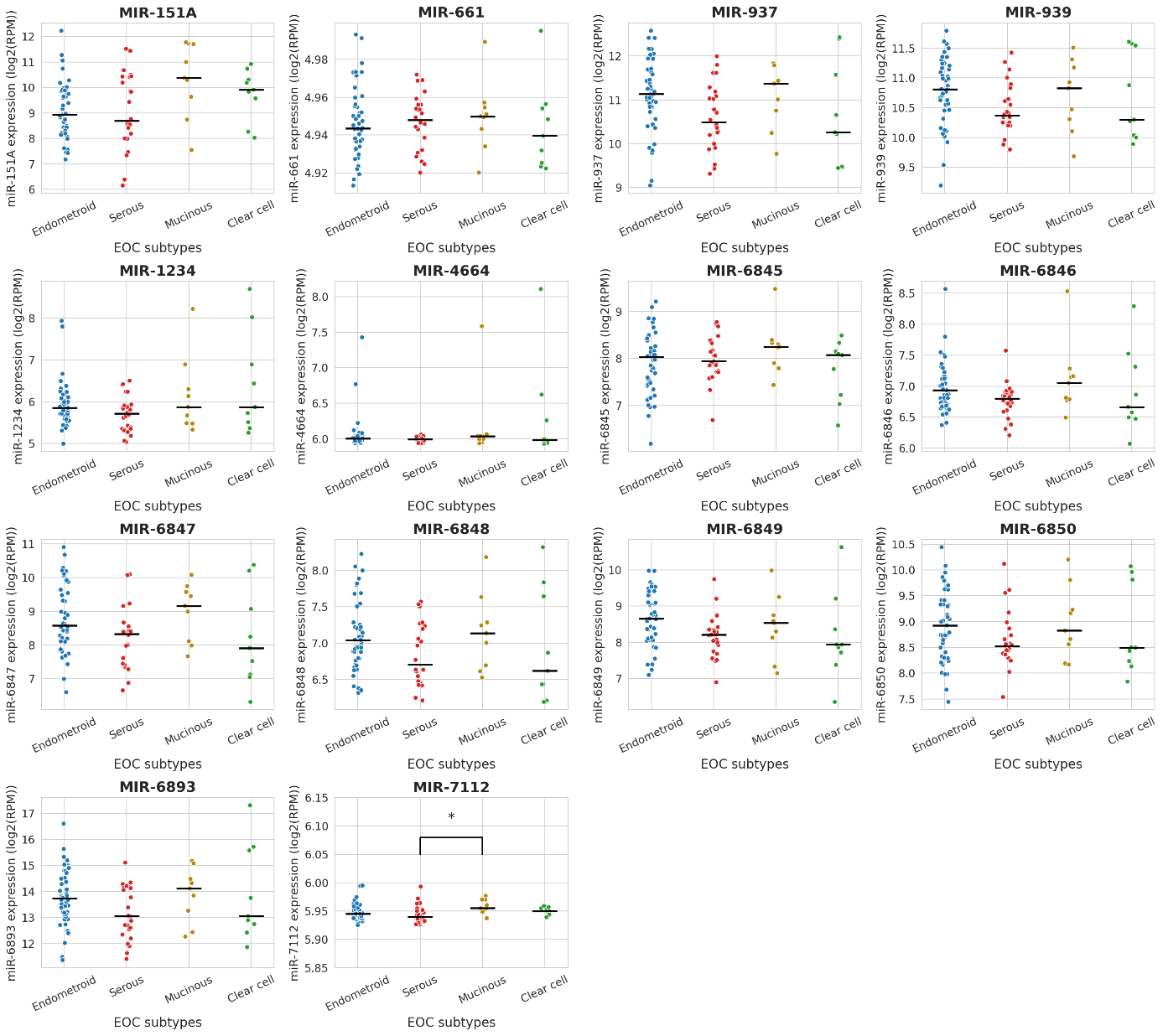

**Fig. S6** **Expression levels of selected 8q24.3-encoded miRNAs in different EOC subtypes.** Each dot represents miRNA expression in patient sample. The horizontal black lines indicating the mean expression. Statistically significant differences of miRNA expression between EOC subtypes are indicated (*p < 0.05).

**Figure S7**

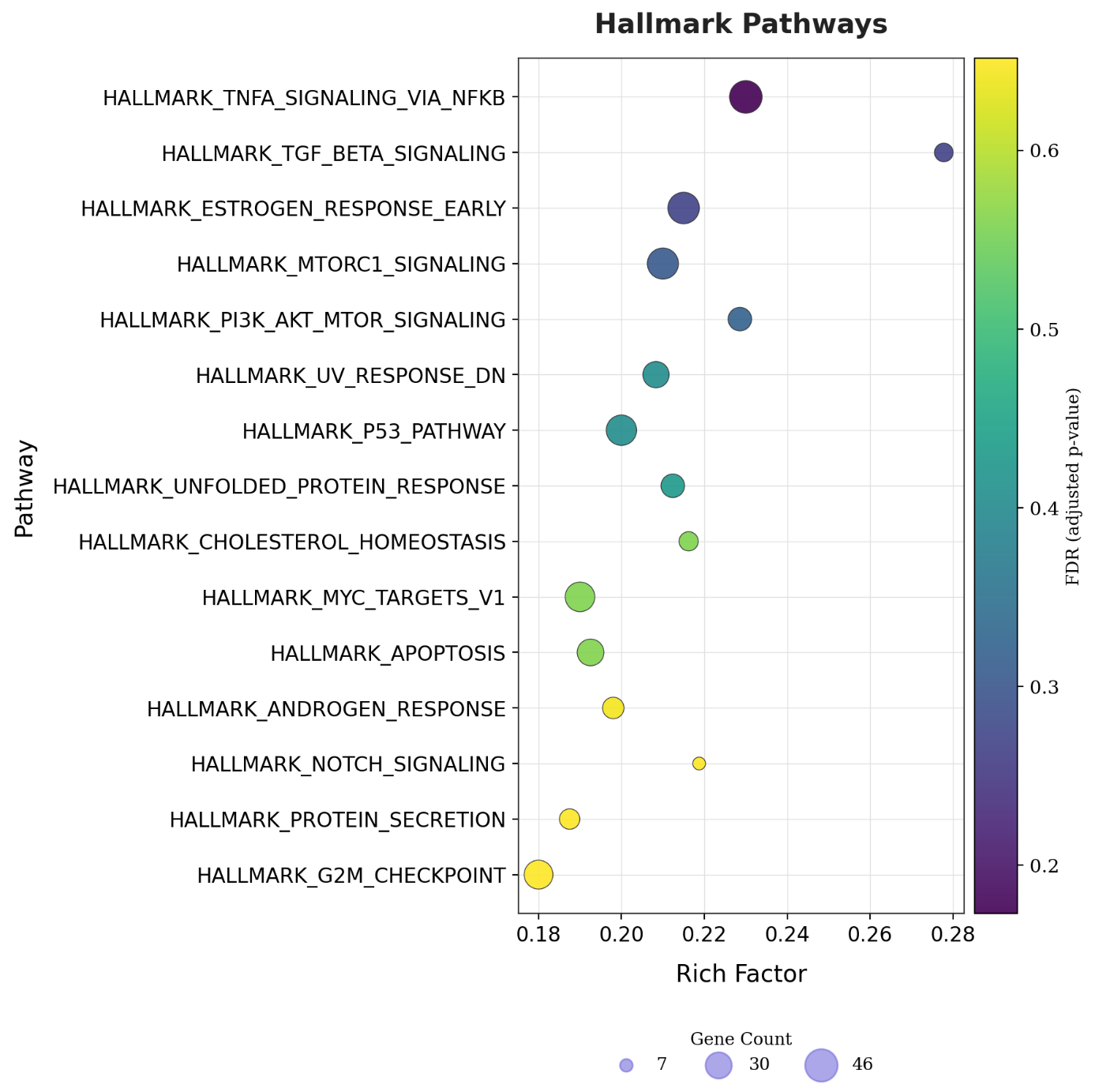

**Fig. S7 Hallmark enrichment analysis of 8q24.3-encoded miRNAs.** Bubble plots display the top 15 enriched Hallmark pathways identified by ORA. Bubble size represents the number of enriched genes (gene count), while bubble color corresponds to the FDR-adjusted p-value.
